# Thalamic Nuclei Functional Controllability Explains Cognition Over and Above Grey and White Matter Structure

**DOI:** 10.64898/2026.05.01.722231

**Authors:** Yuping Yang, Anna Woollams, Marta Czime Litwińczuk, Nelson J Trujillo-Barreto, Nils Muhlert

## Abstract

**Introduction:** The thalamic nuclei play a crucial role in regulating information flow to the cortex and supports diverse cognitive functions. Although previous studies have linked thalamic structural and functional characteristics to cognition, these measures do not fully capture the thalamus’s role in dynamic control, which is essential for complex cognitive processes. Moreover, it remains unclear how these different metrics relate to each other in the way they account for cognition.

**Methods:** T1-weighted MRI, diffusion MRI, resting-state fMRI, and neuropsychological data were obtained from 419 unrelated participants in the Human Connectome Project. We measured grey matter volume, white matter integrity, and functional controllability of each thalamic nucleus to examine their associations with cognitive performance across domains identified through clustering analysis of the neuropsychological data. We also assessed the relationships among these structural and functional metrics and evaluated their individual and combined contributions in capturing covariance with performance in various cognitive domains.

**Results:** Significant correlations were observed between thalamic grey matter volume and white matter integrity; however, thalamic functional controllability showed no significant association with either structural metric. White matter integrity demonstrated the strongest association with sequence working memory and language processing. In contrast, thalamic controllability metrics accounted more for performance in executive function, reasoning and encoding, visuospatial processing, and impulse control, outperforming the combination of grey and white matter structural metrics.

**Conclusion:** This study highlights the critical role of the thalamus from a dynamic control perspective, demonstrating that thalamic structural and functional metrics provide complementary rather than redundant information related to cognitive performance. These findings underscore a promising new direction for understanding the complex and dynamic contributions of the thalamus to human cognition.

## Introduction

The thalamus is a deep brain structure that plays a key role in regulating information flow to cortical regions. Decades of research have established its critical involvement in a wide range of cognitive functions, including working memory (Nadeau, 2008; Roy et al., 2022), executive function (Ferris et al., 2022; Wolff & Halassa, 2024), language processing (Crosson, 2021; Nadeau, 2021), reasoning (Wang, 2024a), cognitive encoding (Escera, 2023; Tankus et al., 2024), visuospatial processing (Rafal & Posner, 1987; Sommer & Wurtz, 2006; Usrey & Alitto, 2015), and impulse control (Guzulaitis & Palmer, 2023a; Zhong et al., 2025). The thalamus comprises multiple nuclei which, despite sharing similar cellular properties, differ in their neurochemical profiles and anatomical and functional connections with cortical and subcortical regions. These distinctions underlie the variability in how individual thalamic nuclei contribute to cognitive functions, as demonstrated by neuropsychological studies investigating the role of specific nuclei in cognitive performance (reviewed in Antonucci et al., 2021).

From a structural perspective, both thalamic grey matter volume and white matter microstructural integrity are associated with cognitive performance. Volumetric studies have reported negative correlations between thalamic volume and cognitive domains such as processing speed and working memory in healthy young and middle-aged adults, even after adjusting for age (Van Der Werf et al., 2001). Similarly, reduced volume in the thalamofrontal projection has been linked to poorer executive function in healthy individuals (Hughes et al., 2012). Beyond normative populations, thalamic atrophy has also been associated with cognitive impairments across various clinical conditions, including multiple sclerosis (Houtchens et al., 2007; Schoonheim et al., 2015), dementia (Biesbroek et al., 2024; Thanissery et al., 2025), stroke (Stebbins et al., 2008), and Alzheimer’s Disease (de Jong et al., 2008; van de Mortel et al., 2021). In addition, diffusion imaging studies have demonstrated associations between thalamic white matter microstructural integrity and cognitive functioning. In healthy individuals, higher fractional anisotropy (FA) within the thalamus has been associated with better executive function (Fama & Sullivan, 2015; Grieve et al., 2007), while higher mean diffusivity (MD) has been linked to lower working memory performance (Piras et al., 2010). Moreover, both lower FA and higher MD in the thalamus have been correlated with slower information processing (Fama & Sullivan, 2015). Similar associations have also been reported in clinical populations, such as decreased thalamic FA and increased thalamic MD in cognitively impaired individuals with multiple sclerosis (Schoonheim et al., 2015); increased thalamic MD with deficits in attention and language processing in Parkinson’s disease (Borlase et al., 2020); as well as decreased thalamic FA and increased thalamic MD with impaired verbal fluency following stroke (Fernández-Andújar et al., 2014).

From a functional perspective, the thalamus is extensively interconnected with multiple regions of the central nervous system, positioning it as a critical hub for modulating brain dynamics and supporting a wide range of cognitive and behavioural functions (Halassa & Sherman, 2019; Sherman, 2007; Shine et al., 2023). Functional connectivity studies have provided evidence linking thalamic connectivity to cognitive performance in both healthy individuals (Goldstone et al., 2018a; K. Li et al., 2022; Wang, 2024b) and clinical populations (M. Li et al., 2020; Schoonheim et al., 2015; Tona et al., 2014; Wu et al., 2022). However, these approaches primarily capture the temporal synchronization of thalamic regions during rest or a specific task state, and therefore are insufficient to fully characterize how the thalamus contributes to dynamic transitions between distinct cognitive processes.

An emerging approach for characterizing how specific brain regions influence dynamic transitions between distinct cognitive processes is network controllability analysis (Lynn & Bassett, 2019; Gu et al., 2015; Deng et al., 2022). Unlike conventional functional connectivity, which assesses the synchronization between regions during a given cognitive state, network controllability offers a framework for identifying the capacity of brain regions to drive transitions between distinct cognitive activity states. This method has proven effective in explaining cognitive performance across both healthy populations (Deng et al., 2022; Deng & Gu, 2020; Gu et al., 2015) and clinical cohorts (Bernhardt et al., 2019; Hahn et al., 2023; Q. Li et al., 2023; Parkes et al., 2021; Yang et al., 2025; Zarkali et al., 2020). However, current research has yet to systematically investigate the network controllability of the thalamus and its subnuclei. It remains unclear whether thalamic controllability correlates with structural profiles and how these factors jointly account for individual differences in cognitive performance.

In this study, we examined thalamic network controllability derived from functional connectivity (hereafter, thalamic functional controllability) and its relationships with thalamic grey matter volume, white matter integrity, and cognition. Our aim was to assess the unique and combined contributions of these metrics to inter-individual variability in cognitive performance. In particular, we evaluated the added value of functional controllability as a novel measure that reflects the capacity of thalamic nuclei to dynamically influence brain-wide activity patterns underpinning cognitive functions. We hypothesised that: (1) controllability metrics of specific thalamic nuclei would be associated with specific cognitive domains; (2) these metrics would be largely independent of grey and white matter measures; (3) they would capture unique variance in cognition rather than overlapping with structural metrics; and (4) combining functional controllability with structural measures would provide a more comprehensive account of cognitive performance than either modality alone.

## Materials and Methods

### Participants

Neuroimaging data were obtained from the publicly available Human Connectome Project (HCP) S1200 dataset,(Van Essen et al., 2013) which includes high-resolution 3T MRI scans acquired using three imaging modalities: structural MRI (T1-weighted; T1w), resting-state functional MRI (rs-fMRI), and high angular resolution diffusion MRI (dMRI). A subset of 419 unrelated participants was selected based on the availability of complete data across all three modalities. The final sample had a mean age of 28.67 years (standard deviation = 3.73) and included 194 males and 225 females. Informed consent was obtained by the HCP for all participants. Data usage for the present study was approved by the HCP and adheres to all relevant ethical regulations for research involving human participants.

### Behavioural data

The present study used a comprehensive set of behavioural data from the HCP dataset (Barch et al., 2013), encompassing cognitive domains assessed through the NIH Toolbox Cognition Battery (http://www.nihtoolbox.org), alongside additional non-Toolbox behavioural measures (Gur et al., n.d.). Specifically, the NIH Toolbox Cognition Battery provided raw cognitive test scores across the following domains: episodic memory, assessed using the Picture Sequence Memory Test (PicSeq); executive function/cognitive flexibility, using the Dimensional Change Card Sort Test (CardSort); executive function/inhibition, using the Flanker Inhibitory Control and Attention Task (Flanker); language/reading decoding, using the Oral Reading Recognition Test (ReadEng); language-vocabulary Comprehension, using the Picture Vocabulary Test (PicVocab); processing speed, using the Pattern Comparison Processing Speed Test (ProcSpeed); and working memory, using the List Sorting Test (ListSort).

Additionally, non-Toolbox measures included: impulse control, assessed using the Delay Discounting Test (DDisc_AUC_200, DDisc_AUC_40K); fluid intelligence, using the Penn Progressive Matrices Test (PMAT24); spatial orientation, using the Variable Short Penn Line Orientation Test (VSPLOT); sustained attention, using the Short Penn Continuous Performance Test (SCPT); and verbal episodic memory, using the Penn Word Memory Test (IWRD). A measure of overall efficiency of each of the tasks was calculated by dividing the median reaction time for correct responses with the accuracy, as described previously (Liesefeld & Janczyk, 2019; Litwińczuk et al., 2022).

### MRI data acquisition

MRI data were acquired using a 3T Siemens Skyra scanner equipped with a 32-channel head coil. High-resolution three-dimensional T1w images were obtained using a 3D MPRAGE sequence. (TR = 2400 ms, TE = 2.14 ms, TI = 1000 ms, flip angle = 8°, matrix size = 320, FOV = 224 mm, BW = 210 Hz per pixel, ES = 7.6 ms, voxel size = 0.7 mm isotropic, 256 slices). A T2-weighted image is acquired using the variable flip angle turbo spin-echo Siemens SPACE sequence (TR = 3200 ms, TE = 565 ms, matrix size = 320, FOV = 224 mm, BW = 744 Hz per pixel, voxel size = 0.7 mm isotropic, 256 slices).(Glasser et al., 2013)

dMRI data was acquired using a 3T Siemens Skyra scanner equipped with a Siemens SC72 gradient coil and a high-performance gradient power supply, providing a maximum gradient amplitude Gmax of 100 mT/m for diffusion encoding (TR = 5520 ms, TE = 89.5 ms, flip angle = 78°, matrix size = 168 × 144 (RO × PE), FOV = 210 × 180 mm (RO × PE), BW = 1488 Hz per pixel, ES = 0.78 ms, voxel size = 1.25 mm isotropic, 111 slices).

rs-fMRI data was acquired in two runs (15 mins each) using a gradient-echo EPI sequence (TR = 720 ms, TE = 33.1 ms, flip angle = 52°, matrix size = 104 × 90 (RO × PE), FOV = 208 × 180 mm (RO × PE), BW = 2290 Hz per pixel, ES = 0.58 ms, voxel size = 2.0 mm isotropic, 72 slices). Oblique axial images were collected with alternating phase-encoding directions: right-to-left (RL) in one run and left-to-right (LR) in the other. During scanning, participants were instructed to keep their eyes open and maintain relaxed fixation on a projected bright crosshair presented on a dark background (Glasser et al., 2013; Uğurbil et al., 2013).

### MRI data preprocessing

The present study used the minimally pre-processed neuroimaging data provided by the HCP. Details of the data preprocessing pipeline are described previously in.(Glasser et al., 2013) Additional preprocessing steps were further performed to the minimal processing pipeline. Briefly, preprocessing pipeline of T1w images included gradient distortion correction, ACPC registration, brain extraction, field map distortion correction, bias field correction, and MNI152 linear and nonlinear registration. A final spatial smoothing step was applied using a Gaussian kernel with a full width at half maximum (FWHM) of 6 mm.

The preprocessing pipeline of dMRI data included intensity normalization, EPI distortion correction, EDDY current and motion correction, gradient nonlinearity correction registration of mean b0 to native volume T1w, transformation of diffusion data, gradient deviation, and gradient directions to structural space.

The preprocessing pipeline of rs-fMRI data included gradient distortion correction, motion correction, field map preprocessing, distortion correction, EPI to T1w registration, one step spline resampling from the original EPI frames to MNI152 space, and intensity normalization. Finally, spatial smoothing was conducted using a Gaussian kernel with a 6-mm FWHM.

### Thalamic grey matter volume

Following preprocessing, a whole-brain grey matter volume map was generated. The subcortical regions were subsequently parcellated into 16 thalamic nuclei - eight per hemisphere - using a standard subcortical parcellation atlas (Figure 1) (Tian et al., 2020). These included (1) two nucleus per hemisphere in the dorsoanterior thalamus, namely medial dorsoanterior thalamus (THA-DAm-rh/ THA-DAm-lh) and lateral dorsoanterior thalamus (THA-DAl-rh/ THA-DAl-lh); (2) three nucleus per hemisphere in the ventroanterior thalamus, namely superior ventroanterior thalamus (THA-VAs-rh/ THA-VAs-lh), anterior inferior ventroanterior thalamus (THA-VAia-rh/ THA-Vaia-lh), and posterior inferior ventroanterior thalamus (THA-VAip-rh/ THA-VAip-lh); (3) one nuclei per hemisphere in the dorsoposterior thalamus (THA-DP-rh/ THA-DP-lh); and (4) two nucleus per hemisphere in the ventroposterior thalamus, namely medial ventroposterior thalamus (THA-VPm-rh/ THA-VPm-lh) and lateral ventroposterior thalamus (THA-VPl-rh/ THA-VPl-lh). The volume of each nucleus was computed by averaging the voxel-wise grey matter volume within its respective anatomical boundary.

**Figure 1.**
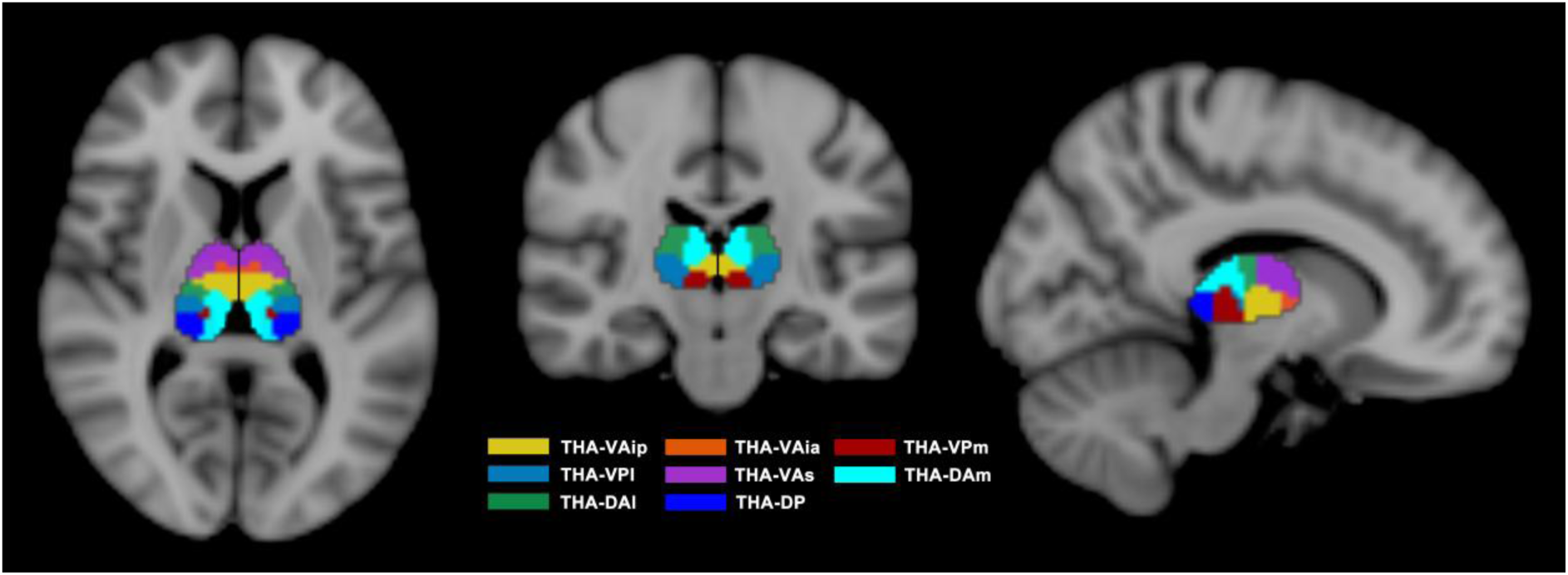
Thalamic nuclei parcellation. The thalamus were parcellated into 16 thalamic nuclei, eight per hemisphere, using a standard subcortical parcellation atlas (Tian et al., 2020). These included two nucleus per hemisphere in the dorsoanterior thalamus, namely medial dorsoanterior thalamus (THA-DAm) and lateral dorsoanterior thalamus (THA-DAl); three nucleus per hemisphere in the ventroanterior thalamus, namely superior ventroanterior thalamus (THA-VAs), anterior inferior ventroanterior thalamus (THA-Vaia) and posterior inferior ventroanterior thalamus (THA-VAip); one nuclei per hemisphere in the dorsoposterior thalamus (THA-DP); and two nucleus per hemisphere in the ventroposterior thalamus, namely medial ventroposterior thalamus (THA-VPm) and lateral ventroposterior thalamus (THA-VPl).

### Thalamic white matter microstructural integrity

The preprocessed dMRI data provided by the HCP were processed using FSL’s *BedpostX* pipeline to model WM fibre orientations and crossing fibres for probabilistic tractography (Glasser et al., 2013). The resulting outputs were then used as inputs for probabilistic tractography with FSL’s *ProbtrackX2*. For each thalamic nucleus, tractography was conducted using the nucleus as a seed mask, with waypoint masks identified by the *Functionnectome* toolbox (Nozais et al., 2023). The cerebellum, brainstem, and ventricles were specified as termination masks and excluded from further analysis to ensure that only cortical and subcortical areas were included, in line with the procedures applied in the volume and functional analyses. Spatial linear and non-linear transformations between standard, diffusion, and individual anatomical spaces were computed using *FLIRT* and *FNIRT*, and subsequently applied during the tractography process in *ProbtrackX*. During tractography, 5000 streamlines were initiated from each voxel, with a step length of 0.5 mm and a maximum of 2000 steps per streamline. Streamline propagation was constrained by a curvature threshold of 0.2 and a minimum volume fraction threshold of 0.01 for fibre orientations. Following these procedures, a probabilistic streamlines map for each thalamic nucleus was obtained. Track density (TD) was then calculated by averaging the probabilistic streamlines passing through each thalamic nucleus. To assess the structural integrity of each thalamic nucleus, whole-brain fractional anisotropy (FA) and mean diffusivity (MD) maps were first generated using DTIFIT. The FA and MD values specific to each thalamic nucleus were then extracted by applying the corresponding track masks derived from tractography.

### Thalamic functional controllability

To examine the functional controllability of each thalamic nucleus, we first constructed a whole brain functional connectivity network using a combined parcellation of 454 ROIs. This included 400 cortical regions defined by the Schaefer atlas, which reflects canonical functional organisation (Schaefer et al., 2018), and 54 subcortical regions derived from a connectivity-gradient-based parcellation (Tian et al., 2020). For each participant, we computed functional connectivity by averaging voxel-wise BOLD signals within each ROI and calculating pairwise Pearson correlations between all ROIs. We then computed controllability metrics for each thalamic nucleus using a linear time-invariant model of brain dynamics, where the system matrix was derived from the normalized graph Laplacian of the functional connectivity network to ensure system stability (Deng et al., 2022; Gu et al., 2015). This framework captures how the thalamic nuclei influence the brain activity transitions. Three controllability metrics were derived: average controllability, modal controllability, and activation energy, as described in our previous work (Yang et al., 2025). Average controllability quantifies the extent to which a node can drive the system into easy-to-reach states with low energy inputs, while modal controllability measures a node’s influence over transitions to difficult-to-reach states with high energy inputs. Activation energy reflects the minimum energy required to engage a node in driving system dynamics. (further details in Supplementary Methods) (Deng et al., 2022; Gu et al., 2015).

### Clustering analysis of cognitive tests

We incorporated both NIH Toolbox cognitive tests (PicSeq, CardSort, Flanker, ReadEng, PicVocab, ProcSpeed, and ListSort) and non-Toolbox cognitive tests (DDisc_AUC_200, DDisc_AUC_40K, PMAT24, VSPLOT, SCPT, and IWRD) into a clustering analysis and categorised them into distinct classes representing commonly used cognitive domains. Specifically, group-level scores were calculated by averaging individual scores for each of the 13 cognitive tests. A total of 10,000 iterations of K-means clustering were then performed on the group-level data to identify robust cognitive domains. The optimal number of clusters (*K*) was selected from the range of 3 to 10 (i.e., 3 < *K* < 10) based on the solution with the highest average silhouette score, which reflects how well each point fits within its assigned cluster compared to others. A final group-level clustering map was produced based on the most stable cluster assignments observed across all 10,000 iterations.

### Statistical analysis

Statistical analyses were performed using MATLAB software version R2022b (MathWorks, Inc). A 95% confidence interval was used for each effect. Bonferroni corrections were performed for multiple comparisons.

### Associations between thalamic imaging metrics and cognitive performance

Correlation analyses were conducted to investigate the associations between thalamic imaging metrics (volume metrics; diffusion metrics: FA, MD, TD; and functional controllability metrics: average controllability, modal controllability, activation energy) and cognitive performance of each of the domains identified through the clustering analysis. Prior to correlation analysis, the distribution of each variable was assessed using the *Lilliefors* test. Partial Pearson’s correlation coefficients were computed for normally distributed variables, while Spearman’s correlations were applied for those not meeting normality assumptions. Age and sex were included as covariates in all analyses involving neuropsychological and imaging metrics.

### Associations among thalamic imaging metrics

To examine whether thalamic imaging metrics derived from different imaging modalities captured overlapping or distinct information relevant to cognitive function, correlation analyses were also performed among thalamic volume metrics, diffusion metrics (FA, MD, TD), and functional controllability metrics (average controllability, modal controllability, activation energy) metrics. As above, Lilliefors tests were used to assess normality, followed by Pearson’s or Spearman’s correlations as appropriate. Age and sex were included as covariates in all analyses involving imaging metrics.

### Covariance between thalamic imaging metrics and cognitive performance

To further examine the covariance between thalamic imaging metrics and cognitive performance, we conducted sparse canonical correlation analysis (CCA) (Mackey, 2008; Uurtio et al., 2019) under various combinations of imaging modalities. Specifically, we explored whether individual modalities or their combinations were more effective in capturing covariance with performance across specific cognitive domains and overall cognitive function. The combinations of imaging metrics included: 1) Single modality: volume metrics only, diffusion metrics (FA, MD, TD) only, or functional controllability metrics (average controllability, modal controllability, activation energy) only; 2) Dual modality: volume + diffusion, volume + functional, or diffusion + functional; and 3) Triple modality: volume, diffusion, and functional metrics combined. All variables were z-scored prior to analysis. The CCA was implemented using an alternating projected gradient algorithm to identify sparse canonical coefficients between imaging and cognitive measures. Regularisation parameters were optimised via 10-fold cross-validation using grid search. Initialisation was performed using left/right singular vectors and random vectors derived from the cross-covariance matrix. The optimal model was selected based on the average test set correlation.(Mackey, 2008; Uurtio et al., 2019)

## Results

### Cognitive domains

All cognitive tests were clustered into six domains: (1) Sequential working memory, including PicSeq and ListSort; (2) Executive function, including CardSort, Flanker, and ProcSpeed; (3) Language processing, including ReadEng and PicVocab; (4) Reasoning and encoding, including PMAT24, IWRD, and SCPT; (5) Visuospatial processing, including VSPLOT; and (6) Impulse control, including DDisc_AUC_200 and DDisc_AUC_40K. Figure 2 presents the group-level clustering map derived from the probability of cluster assignments across all 10,000 iterations.

**Figure 2.**
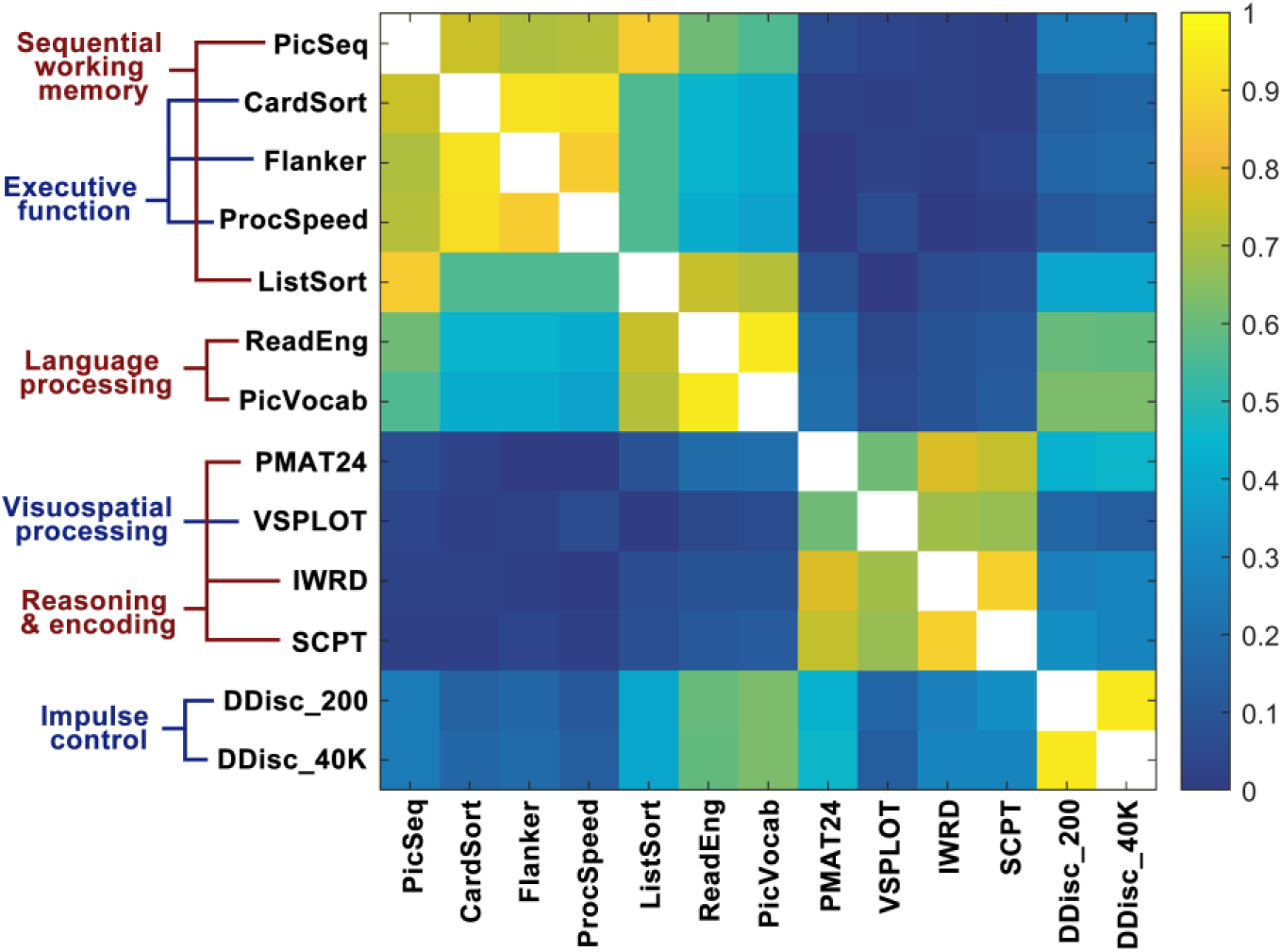
The group-level clustering map of cognitive domains. All cognitive tests were clustered into six domains: (1) Sequential working memory, including PicSeq and ListSort; (2) Executive function, including CardSort, Flanker, and ProcSpeed; (3) Language processing, including ReadEng and PicVocab; (4) Reasoning and encoding, including PMAT24, IWRD, and SCPT; (5) Visuospatial processing, including VSPLOT; and (6) Impulse control, including DDisc_AUC_200 and DDisc_AUC_40K.

### Associations between thalamic imaging metrics and cognitive performance

#### Associations between thalamic volume metrics and cognitive performance

As shown in Figure 3, significant correlations were found between performance in sequential working memory and thalamic volume in THA-DP-rh (rho = 0.143, p = 0.002), THA-VAia-lh (rho = 0.142, p = 0.003), and THA-DP-lh (rho = 0.144, p = 0.002).

**Figure 3.**
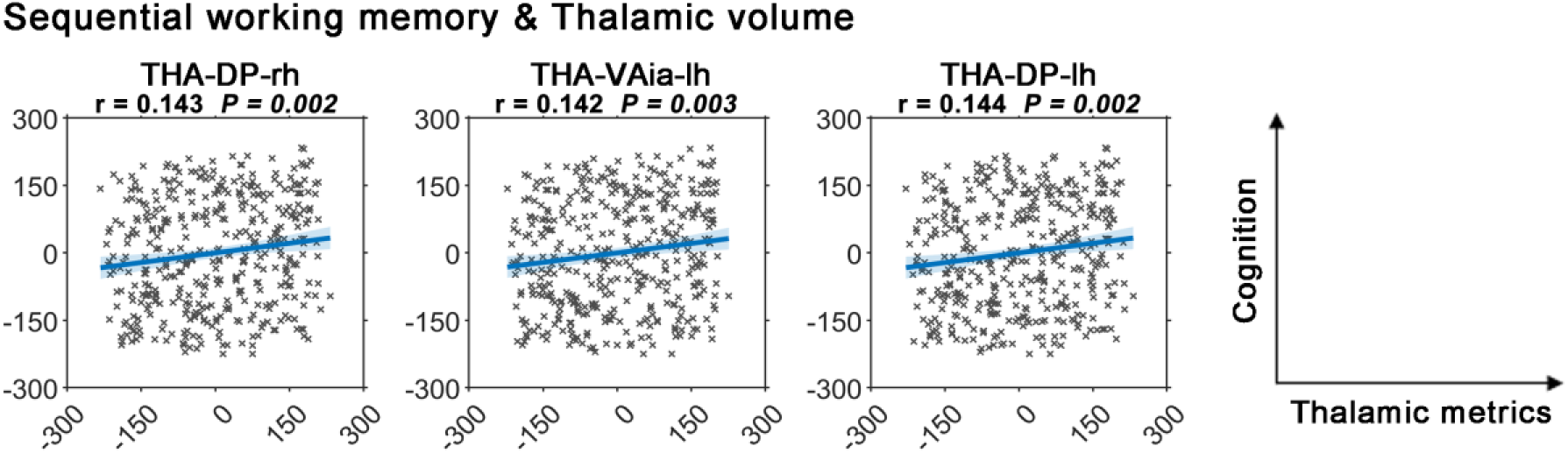
Associations between thalamic volume metrics and cognitive performance. significant correlations were found between performance in sequential working memory and thalamic volume in THA-DP-rh, THA-VAia-lh, and THA-DP-lh.

#### Associations between thalamic diffusion metrics and cognitive performance

As shown in Figure 4, significant correlations were found between performance in language processing and thalamic MD in all thalamic nuclei: THA-VAip-rh (rho = 0.227, p < 0.001), THA-VAia-rh (rho = 0.231, p < 0.001), THA-VPm-rh (rho = 0.230, p < 0.001), THA-VPl-rh (rho = 0.230, p < 0.001), THA-VAs-rh (rho = 0.228, p < 0.001), THA-DAm-rh (rho = 0.230, p < 0.001), THA-DAl-rh (rho = 0.228, p < 0.001), THA-DP-rh (rho = 0.231, p < 0.001), THA-VAip-lh (rho = 0.227, p < 0.001), THA-VAia-lh (rho = 0.221, p < 0.001), THA-VPm-lh (rho = 0.229, p < 0.001), THA-VPl-lh (rho = 0.234, p < 0.001), THA-VAs-lh (rho = 0.228, p < 0.001), THA-DAm-lh (rho = 0.229, p < 0.001), THA-DAl-lh (rho = 0.230, p < 0.001), and THA-DP-lh (rho = 0.233, p < 0.001) (Figure 4A); besides, significant correlations were found between performance in sequential working memory and thalamic MD in THA-VAip-rh (rho = 0.142, p = 0.003), THA-VPm-rh (rho = 0.142, p = 0.003), THA-VPl-rh (rho = 0.143, p = 0.002), THA-VAs-rh (rho = 0.144, p = 0.002), THA-DAl-rh (rho = 0.144, p = 0.002), THA-VAip-lh (rho = 0.141, p = 0.003), THA-VPl-lh (rho = 0.148, p = 0.002), THA-VAs-lh (rho = 0.143, p = 0.002), THA-DAl-lh (rho = 0.146, p = 0.002), and THA-DP-lh (rho = 0.142, p = 0.003) (Figure 4B); additionally, significant correlations were found between performance in visuospatial processing and thalamic FA in THA-VAia-rh (rho = -0.143, p = 0.002), THA-VPl-rh (rho = -0.147, p = 0.002), and THA-DP-lh (rho = -0.141, p = 0.003) (Figure 4C).

**Figure 4.**
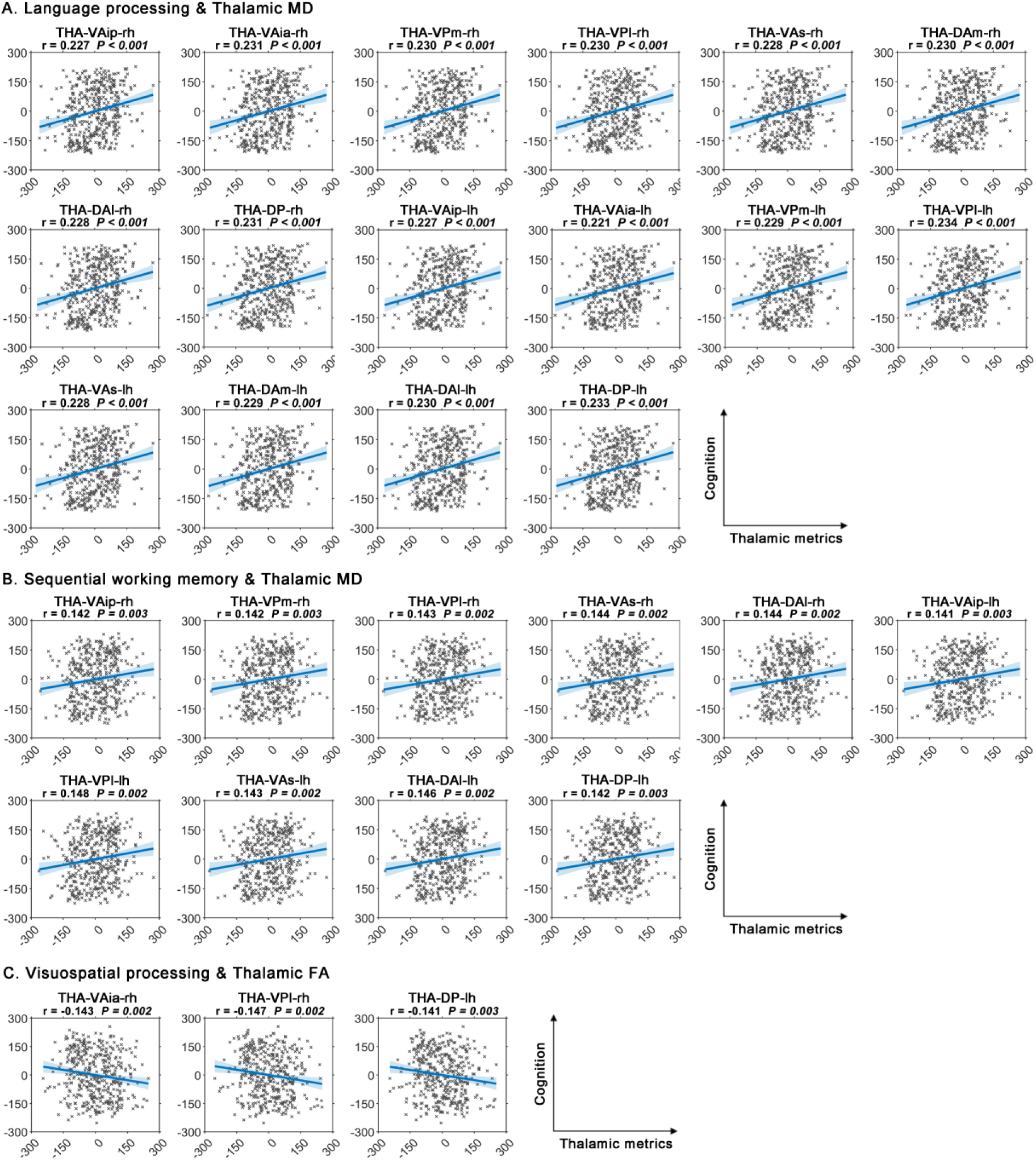
Associations between thalamic diffusion metrics and cognitive performance. **(4A)** Significant correlations were found between performance in language processing and thalamic MD in all thalamic nuclei. **(4B)** Significant correlations were found between performance in sequential working memory and thalamic MD in THA-VAip-rh, THA-VPm-rh, THA-VPl-rh, THA-VAs-rh, THA-DAl-rh, THA-VAip-lh, THA-VPl-lh, THA-VAs-lh, THA-DAl-lh, and THA-DP-lh. **(4C)** Significant correlations were found between performance in visuospatial processing and thalamic FA in THA-VAia-rh, THA-VPl-rh, and THA-DP-lh.

#### Associations between thalamic controllability metrics and cognitive performance

As shown in Figure 5, significant correlations were found between performance in language processing and thalamic average controllability in THA-VAia-rh (rho = 0.140, p = 0.003), THA-DAl-rh (rho = 0.142, p = 0.003), and THA-VAia-lh (rho = 0.146, p = 0.002) (Figure 5A); besides, significant correlations were found between performance in language processing and thalamic modal controllability in THA-VAia-rh (rho = -0.141, p = 0.003), THA-DAl-rh (rho = - 0.142, p = 0.003), and THA-VAia-lh (rho = -0.147, p = 0.002) (Figure 5B); additionally, significant correlations were found between performance in language processing and thalamic activation energy in THA-VAia-rh (rho = -0.142, p = 0.003), THA-DAl-rh (rho = - 0.142, p = 0.003), and THA-VAia-lh (rho = -0.148, p = 0.002) (Figure 5C).

**Figure 5.**
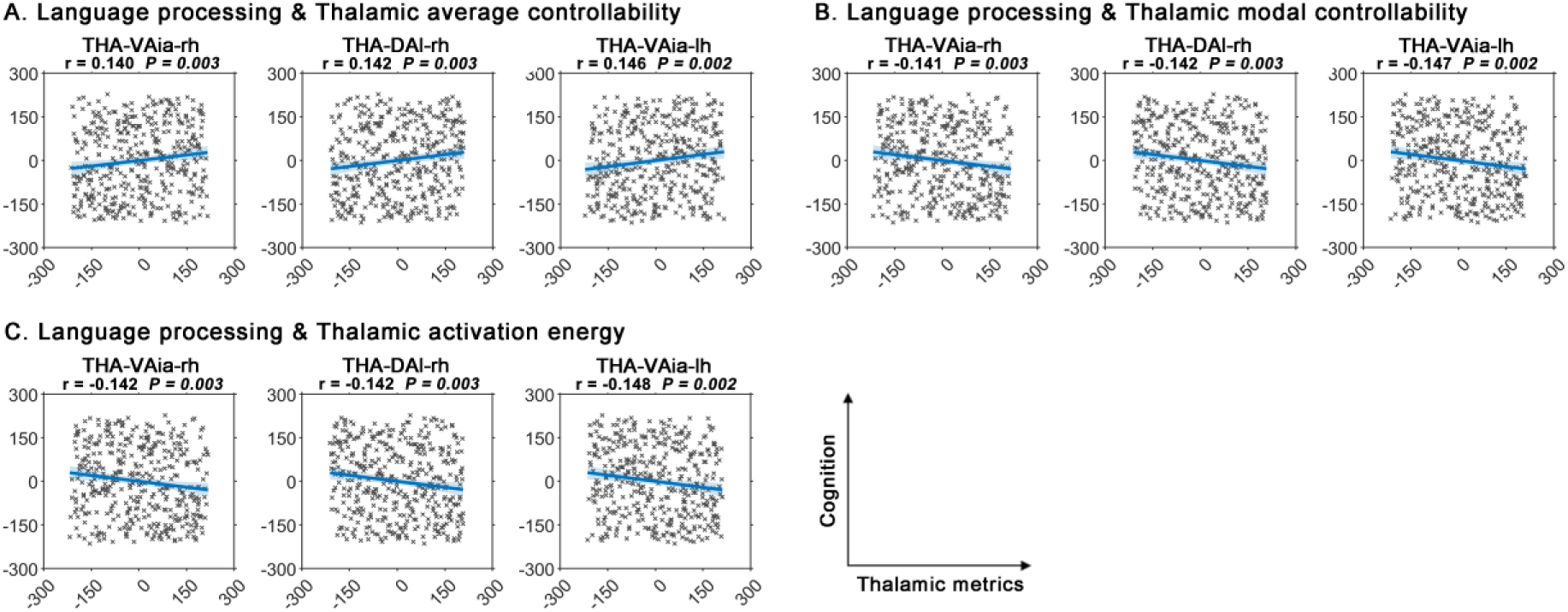
Associations between thalamic controllability metrics and cognitive performance. **(5A)** Significant correlations were found between performance in language processing and thalamic average controllability in THA-VAia-rh, THA-DAl-rh, and THA-VAia-lh. **(5B)** Significant correlations were found between performance in language processing and thalamic modal controllability in THA-VAia-rh, THA-DAl-rh, and THA-VAia-lh. **(5C)** Significant correlations were found between performance in language processing and thalamic activation energy in THA-VAia-rh, THA-DAl-rh, and THA-VAia-lh.

### Associations among thalamic imaging metrics

Significant correlations were found between volume and diffusion metrics (Figure 6). Specifically, significant positive correlations were found between thalamic volume and thalamic FA in all thalamic nuclei: THA-VAip-rh (rho = 0.240, *p* < 0.001), THA-VAia-rh (rho = 0.236, *p* < 0.001), THA-VPm-rh (rho = 0.246, *p* < 0.001), THA-VPl-rh (rho = 0.232, *p* < 0.001), THA-VAs-rh (rho = 0.232, *p* < 0.001), THA-DAm-rh (rho = 0.172, *p* < 0.001), THA-DAl-rh (rho = 0.219, *p* < 0.001), THA-DP-rh (rho = 0.241, *p* < 0.001), THA-VAip-lh (rho = 0.238, *p* < 0.001), THA-VAia-lh (rho = 0.230, *p* < 0.001), THA-VPm-lh (rho = 0.238, *p* < 0.001), THA-VPl-lh (rho = 0.225, *p* < 0.001), THA-VAs-lh (rho = 0.239, *p* < 0.001), THA-DAm-lh (rho = 0.163, *p* < 0.001), THA-DAl-lh (rho = 0.223, *p* < 0.001), and THA-DP-lh (rho = 0.241, *p* < 0.001 (Figure 6A). Besides, significant positive correlations were found between thalamic volume and thalamic MD in all thalamic nuclei: THA-VAip-rh (rho = 0.306, *p* < 0.001), THA-VAia-rh (rho = 0.289, *p* < 0.001), THA-VPm-rh (rho = 0.308, *p* < 0.001), THA-VPl-rh (rho = 0.319, *p* < 0.001), THA-VAs-rh (rho = 0.294, *p* < 0.001), THA-DAm-rh (rho = 0.211, *p* < 0.001), THA-DAl-rh (rho = 0.302, *p* < 0.001), THA-DP-rh (rho = 0.284, *p* < 0.001), THA-VAip-lh (rho = 0.294, *p* < 0.001), THA-VAia-lh (rho = 0.264, *p* < 0.001), THA-VPm-lh (rho = 0.286, *p* < 0.001), THA-VPl-lh (rho = 0.311, *p* < 0.001), THA-VAs-lh (rho = 0.281, *p* < 0.001), THA-DAm-lh (rho = 0.181, *p* < 0.001), THA-DAl-lh (rho = 0.289, *p* < 0.001), and THA-DP-lh (rho = 0.281, *p* < 0.001) (Figure 6B). Additionally, significant correlations were found between thalamic volume and thalamic TD in THA-VAia-rh (rho = -0.156, *p* = 0.001) and THA-DAm-lh (rho = 0.216, *p* < 0.001) (Figure 6C). No significant correlations were found between controllability metrics and either volume or diffusion metrics (*p* > 0.05).

**Figure 6.**
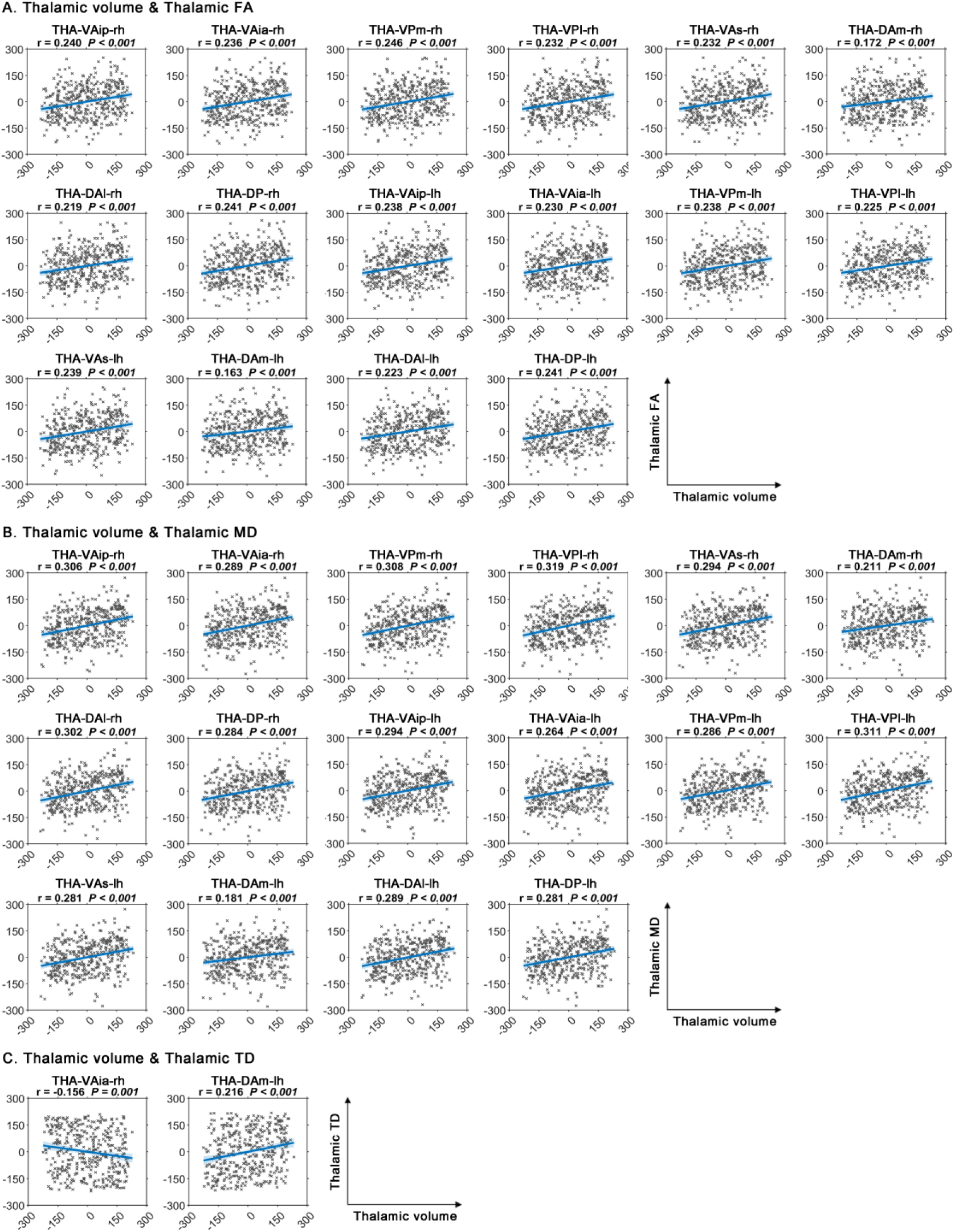
Associations among thalamic imaging metrics. Significant correlations were found between volume and diffusion metrics, but no significant correlations were found between controllability metrics and either volume or diffusion metrics. **(6A)** Significant positive correlations were found between thalamic volume and thalamic FA in all thalamic nuclei. **(6B)** Significant positive correlations were found between thalamic volume and thalamic MD in all thalamic nuclei. **(6C)** Significant correlations were found between thalamic volume and thalamic TD in THA-VAia-rh and THA-DAm-lh.

### Covariance between thalamic imaging metrics and cognitive performance

#### Sequential working memory

Diffusion metrics exhibited the highest covariance with performance in sequential working memory (Figure 7A, Supplementary Table 1). Specifically, the strongest associations were observed with the triple-modality model combining controllability, diffusion, and volume metrics (R = 0.510). Among dual-modality models, the combination of controllability and diffusion metrics yielded the highest association (R = 0.489), followed by controllability and volume (R = 0.407), and diffusion and volume (R = 0.396). For single-modality models, diffusion metrics showed the strongest associations (R = 0.353), followed by controllability (R = 0.347), and volume metrics (R = 0.296).

**Figure 7.**
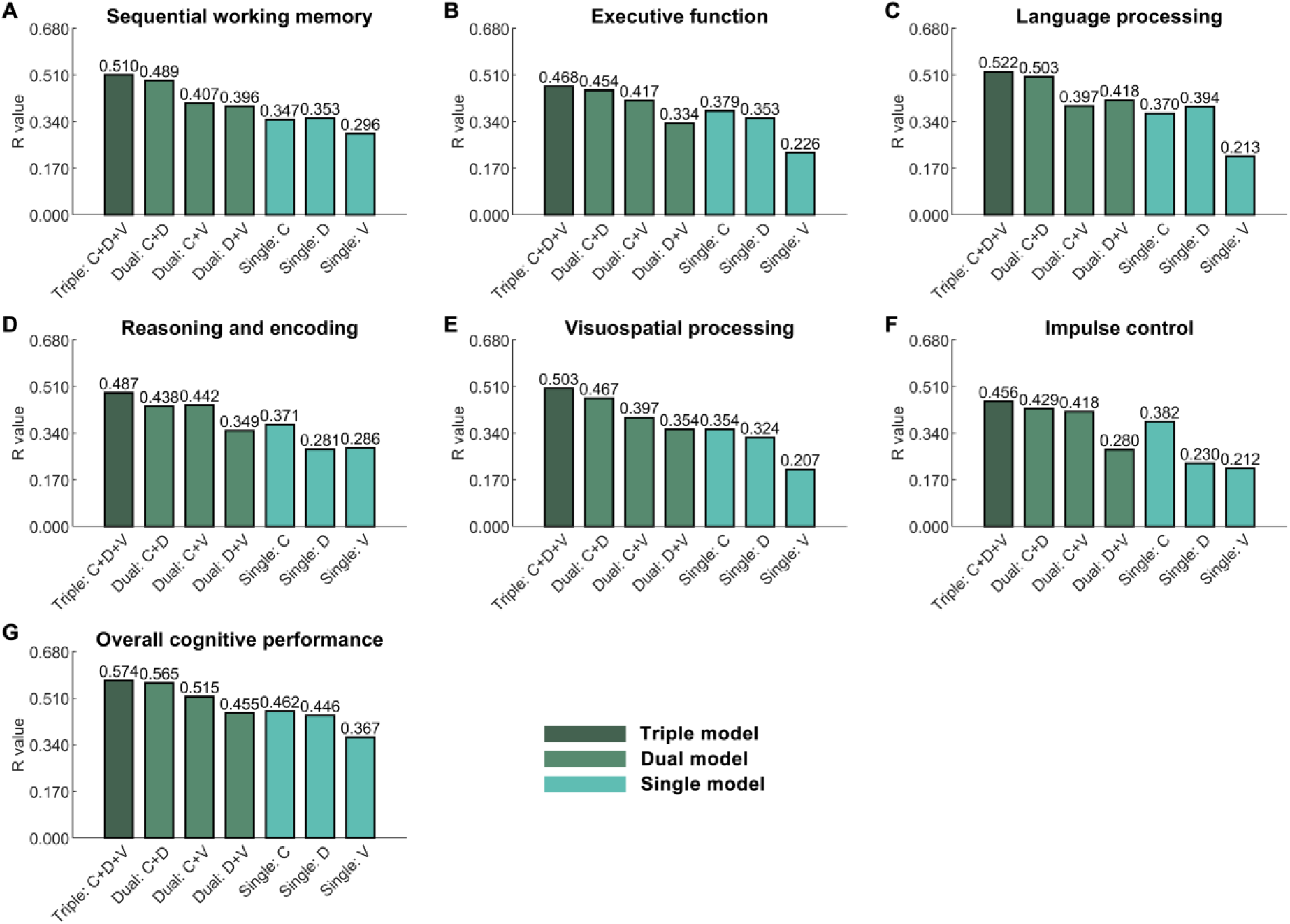
Covariance between thalamic imaging metrics and cognitive performance. **(8A)** Diffusion metrics exhibited the highest covariance with performance in sequential working memory. **(7B)** Controllability metrics exhibited the highest covariance with performance in executive function. **(7C)** Diffusion metrics exhibited the highest covariance with performance in language processing. **(7D)** Controllability metrics exhibited the highest covariance with performance in reasoning and encoding. **(7E)** Controllability metrics exhibited the highest covariance with performance in visuospatial processing. **(7F)** Controllability metrics exhibited the highest covariance with performance in impulse control. **(7G)** Controllability metrics exhibited the highest covariance with overall cognitive performance.

#### Executive function

Controllability metrics exhibited the highest covariance with performance in executive function (Figure 7B, Supplementary Table 2). Specifically, the strongest associations were observed with the triple-modality model combining controllability, diffusion, and volume metrics (R = 0.468). Among dual-modality models, the combination of controllability and diffusion metrics yielded the highest association (R = 0.454), followed by controllability and volume (R = 0.417), and diffusion and volume (R = 0.334). For single-modality models, controllability metrics showed the strongest associations (R = 0.379), followed by diffusion (R = 0.353), and volume metrics (R = 0.226). Notably, controllability metrics on their own (R = 0.379) captured covariance with performance in executive function more effectively than diffusion metrics (R = 0.353), volume metrics (R = 0.226), or their combination (R = 0.334).

#### Language processing

Diffusion metrics exhibited the highest covariance with performance in language processing (Figure 7C, Supplementary Table 3). Specifically, the strongest associations were observed with the triple-modality model combining controllability, diffusion, and volume metrics (R = 0.522). Among dual-modality models, the combination of controllability and diffusion metrics yielded the highest association (R = 0.503), followed by diffusion and volume (R = 0.418), and controllability and volume (R = 0.397). For single-modality models, diffusion metrics showed the strongest associations (R = 0.394), followed by controllability (R = 0.370), and volume metrics (R = 0.213).

#### Reasoning and encoding

Controllability metrics exhibited the highest covariance with performance in reasoning and encoding (Figure 7D, Supplementary Table 4). Specifically, the strongest associations were observed with the triple-modality model combining controllability, diffusion, and volume metrics (R = 0.487). Among dual-modality models, the combination of controllability and volume metrics yielded the highest association (R = 0.442), followed by controllability and diffusion (R = 0.438), and diffusion and volume (R = 0.349). For single-modality models, controllability metrics showed the strongest associations (R = 0.371), followed by volume (R = 0.286), and diffusion metrics (R = 0.281). Again, controllability metrics on their own (R = 0.371) captured covariance with performance in reasoning and encoding more effectively than diffusion metrics (R = 0.281), volume metrics (R = 0.286), or their combination (R = 0.349).

#### Visuospatial processing

Controllability metrics exhibited the highest covariance with performance in visuospatial processing (Figure 7E, Supplementary Table 5). Specifically, the strongest associations were observed with the triple-modality model combining controllability, diffusion, and volume metrics (R = 0.503). Among dual-modality models, the combination of controllability and diffusion metrics yielded the highest association (R = 0.467), followed by controllability and volume (R = 0.397), and diffusion and volume (R = 0.354). For single-modality models, controllability metrics showed the strongest associations (R = 0.354), followed by diffusion (R = 0.324), and volume metrics (R = 0.207). Similarly, controllability metrics on their own (R = 0.354) captured covariance with performance in visuospatial processing more effectively than diffusion metrics (R = 0.324), volume metrics (R = 0.207), while performing equally well as the combined diffusion and volume metrics (R = 0.354).

#### Impulse control

Controllability metrics exhibited the highest covariance with performance in impulse control (Figure 7F, Supplementary Table 6). Specifically, the strongest associations were observed with the triple-modality model combining controllability, diffusion, and volume metrics (R = 0.456). Among dual-modality models, the combination of controllability and diffusion metrics yielded the highest association (R = 0.429), followed by controllability and volume (R = 0.418), and diffusion and volume (R = 0.280). For single-modality models, controllability metrics showed the strongest associations (R = 0.382), followed by diffusion (R = 0.230), and volume metrics (R = 0.212). Repeatedly, controllability metrics on their own (R = 0.382) captured covariance with performance in impulse control more effectively than diffusion metrics (R = 0.230), volume metrics (R = 0.212), or their combination (R = 0.280).

#### Overall cognitive performance

Controllability metrics exhibited the highest covariance with overall cognitive performance (Figure 7G, Supplementary Table 7). Specifically, the strongest associations were observed with the triple-modality model combining controllability, diffusion, and volume metrics (R = 0.574). Among dual-modality models, the combination of controllability and diffusion metrics yielded the highest association (R = 0.565), followed by controllability and volume (R = 0.515), and diffusion and volume (R = 0.455). For single-modality models, controllability metrics showed the strongest associations (R = 0.462), followed by diffusion (R = 0.446), and volume metrics (R = 0.367). Again, controllability metrics on their own (R = 0.462) captured covariance with performance in overall cognition more effectively than diffusion metrics (R = 0.446), volume metrics (R = 0.367), or their combination (R = 0.455).

## Discussion

This study characterised the contribution of distinct thalamic nuclei to cognitive performance across major cognitive domains, drawing on thalamic metrics derived from grey matter volume, white matter microstructural integrity, and functional controllability. Our findings demonstrate that functional controllability in specific thalamic nuclei accounts for unique variance not captured by structural measures alone, and exhibited higher covariance with cognitive performance, particularly in executive function, reasoning and encoding, visuospatial processing, and impulse control.

Our findings showed that thalamic controllability metrics exhibited highest covariance with performance in executive function, reasoning and encoding, visuospatial processing, and impulse control, outperforming diffusion metrics, volume metrics, or even their combination. This underscores the unique contribution of thalamic controllability metrics to explaining cognitive performance, capturing aspects not accounted for by grey matter volume or white matter microstructural integrity. Previous research has also emphasized the importance of thalamic functional properties in cognition. For instance, thalamic lesion sites linked to more severe executive dysfunction exhibit stronger functional connectivity with the anterior cingulate cortex, dorsomedial prefrontal cortex, and the frontoparietal network—regions whose impairment is known to contribute to deficits in executive function (Hwang et al., 2020). Also, research has shown that cognitive reasoning processes are associated with increased activation of different thalamic nuclei and their functional connections with specific cortical regions, with thalamic activation and connectivity further increasing as the relational complexity of reasoning tasks rises (Wang, 2024c). Besides, increased thalamo-hippocampal functional connectivity has been observed in normal ageing, with the largest increases seen in individuals who developed visuospatial processing impairments. (Goldstone et al., 2018b). Additionally, thalamocortical pathways have been shown to exhibit increased neural activity during impulsive behaviour, suggesting a potential role for the thalamus in driving the cortical activities underlying impulsivity (Guzulaitis & Palmer, 2023b). These studies have primarily focused on thalamic functional activation or connectivity, which reflects brain activity in a single state. However, such approaches fail to capture how the thalamus influences cortical regions during transitions between different brain states, which are essential for the flexibility and efficiency of higher-order cognitive functions (Deng et al., 2022). Addressing this gap, our study demonstrates that thalamic functional controllability captures covariance with performance in these cognitive domains more effectively than structural measures alone, offering novel insights into the thalamus’s dynamic control over the cortex and its influence on cognitive performance. These findings point to a promising new direction for understanding the complex and dynamic role of the thalamus in human cognition.

Our results demonstrate that thalamic structural metrics contribute differentially across cognitive domains, particularly in sequential working memory and language processing. Specifically, we found significant associations between sequential working memory performance and both mean diffusivity and volume in several anterior thalamic nuclei. Among the three modalities examined, thalamic diffusion metrics showed the strongest association with individual differences in sequential working memory, as revealed by the CCA models. These findings highlight the involvement of anterior thalamic structures in supporting sequential working memory, consistent with previous animal studies showing that excitotoxic lesions to the anterior thalamic nuclei impair performance in sequential memory tasks (Wolff et al., 2006). Extending this line of evidence to humans, our findings offer converging support for the critical role of anterior thalamic structural integrity in sequential working memory. In addition, we observed significant associations between language processing performance and thalamic mean diffusivity across the whole thalamus. The thalamus has long been recognised for its involvement in language processing. Anatomically, the thalamus forms part of a thalamocortical loop that includes structural connections to Broca’s area, supporting semantic, phonological, and syntactic processing (Ford et al., 2013). Neuroimaging studies have also linked thalamic lesions to thalamic aphasia, typically characterised by anomia, verbal paraphasias, reduced fluency and verbal output, and impaired grammar, while comprehension remains relatively preserved (reviewed in (Barbas et al., 2013; Bulut & Hagoort, 2024). More recently, structural fibre disconnections caused by thalamic lesions have been associated with language impairments in aphasia, further underscoring the role of thalamic fibre integrity in language (Stockert et al., 2025).

Importantly, our findings suggest that the thalamus contributes to language processing not only through its structural properties but also via functional controllability. Moreover, the combined use of thalamic structural and functional measures accounts more for language performance, indicating that these modalities provide complementary rather than redundant information. This observation aligns with previous research emphasising the role of thalamic functional properties in language processing. For instance, several studies have reported associations between thalamic functional connectivity and language performance (Klostermann & Ehlen, 2013; Llano, 2013). One review posits that, rather than engaging in linguistic operations directly, the thalamus functions as a central monitor of language-related cortical activity (Klostermann & Ehlen, 2013). This view is consistent with our findings, which revealed a significant association between thalamic functional controllability and language performance, underscoring the thalamus’s influence on cortical areas in language-related activity through functional network dynamics. Collectively, these results highlight the multifaceted role of the thalamus in language processing and further support the value of integrating structural and functional imaging metrics to better characterise its contribution to cognition.

Our study demonstrates that structural and functional properties of the thalamic nuclei contribute differently to cognitive performance across domains. This finding supports the broader notion that structural and functional properties of the human brain can play both convergent and divergent roles in supporting cognitive functioning (Park & Friston, 2013). The human brain is a complex neural system whose functional organisation is grounded in its underlying structural architecture. While anatomical architecture constrains possible patterns of neural activity, the brain has developed highly dynamic and flexible functionality that emerge from, but are not strictly determined by, its physical structure. This diverse functionality enables the relatively fixed structural scaffold to support diverse and adaptive patterns of neural activity, underpinning a wide range of cognitive functions (Park & Friston, 2013). Recent research has validated the interplay between structural and functional properties and their distinct roles in explaining cognition. For example, functional connectivity has been shown to be one of the strongest predictors of cognition, yet among the weakest features for identifying individuals; while structural connectivity, on the contrary, has proven to be one of the strongest features for individual identification, but a relatively weak predictor of cognition (Mansour L et al., 2021). Another recent study showed that structural connectivity, functional connectivity, and their combination contribute differently to predicting performance in executive function, language, encoding, and sequence processing (Litwińczuk et al., 2022). Together, this body of evidence suggests that brain structure and function support cognition in complementary ways, each contributing distinct aspects. In line with this, our findings revealed that thalamic structural and functional properties were not significantly correlated, and that integrating both modalities exhibited higher covariance with cognitive performance. These results demonstrate that, similar to whole-brain findings, grey matter volume, white matter microstructural integrity, and functional controllability within the thalamic nuclei each offer complementary insights into the neural basis of cognitive performance. This underscores the critical role of the thalamic nuclei in supporting cognitive functions and highlights the importance of future research that combines structural and functional approaches to achieve a more comprehensive understanding of cognition.

Our study has several limitations. First, it focused exclusively on structural and resting-state functional metrics, whereas functional metrics derived from cognitive task-based fMRI might offer additional insights into cognitive performance. Second, this study only used a single thalamic parcellation scheme; however, alternative parcellation approaches may yield different results. Therefore, future research should undertake more comprehensive analyses incorporating multiple thalamic parcellation frameworks to validate and extend these findings. Third, this study examined only linear relationships between thalamic imaging metrics and cognitive performance. Future research should perform more advanced statistical methods to capture potential nonlinear and complex interactions, thereby providing a more nuanced understanding of the thalamus’s role in cognition.

## Conclusion

In summary, this study provides comprehensive evidence that both structural and functional properties of the thalamic nuclei play distinct and complementary roles in supporting cognitive performance across multiple domains. Importantly, our findings highlight the potential of thalamic functional controllability over thalamic grey and white matter structural measures in capturing covariance in cognition, particularly in executive function, reasoning, visuospatial processing, and impulse control. By revealing how the thalamus dynamically influences cortical activity during brain state transitions, this work advances our understanding of the thalamus as a key hub in the neural architecture underlying human cognition. These insights not only deepen our knowledge of thalamic contributions to cognitive function but also pave the way for future research integrating multimodal neuroimaging approaches to unravel the complex mechanisms of cognitive processing and dysfunction.

## Data and Code Availability

Data is openly available as part of the Human Connectome Project S1200 dataset (https://www.humanconnectome.org/study/hcp-young-adult/document/1200-subjects-data-release).

## Author Contributions

Y.Y., N.T.B., and N.M. contributed to the conception and design of the study. Y.Y. and M.C.L. contributed to the acquisition of data. Y.Y., A.W., N.T.B., and N.M. contributed to analysis or interpretation of data. Y.Y., A.W., N.T.B., and N.M. contributed to drafting/revision of the manuscript.

## Declaration of Competing Interests

Y.Y., A.W., M.C.L. and N.M. report no disclosures. N.T.B. reports research grants from the Medical Research Council UK (MR/X005267/1) during the conduct of the study.

## Acknowledgements

We extend our gratitude to all our colleagues from the MISC and SPiN labs for their kind support during this study.

## Supplementary Material

Supplementary materials are available online.

