## Supplementary Materials for "Thalamic Nuclei Functional Controllability Explains Cognition Over and Above Grey and White Matter Structure"

**Supplementary Table 1. Covariance between thalamic imaging metrics and performance in sequential working memory**

| Modality | Thalamic imaging metrics | R |
|---|---|---|
| Triple-modality model | Controllability + Diffusion + Volume | 0.510 |
| Dual-modality model | Controllability + Diffusion | 0.489 |
| | Controllability + Volume | 0.407 |
| | Diffusion + Volume | 0.396 |
| Single-modality model | Controllability | 0.347 |
| | Diffusion | 0.353 |
| | Volume | 0.296 |

R values are generated using sparse canonical correlation analysis.

**Supplementary Table 2. Covariance between thalamic imaging metrics and performance in executive function**

| Modality | Thalamic imaging metrics | R |
|---|---|---|
| Triple-modality model | Controllability + Diffusion + Volume | 0.468 |
| Dual-modality model | Controllability + Diffusion | 0.454 |
| | Controllability + Volume | 0.417 |
| | Diffusion + Volume | 0.334 |
| Single-modality model | Controllability | 0.379 |
| | Diffusion | 0.353 |
| | Volume | 0.226 |

R values are generated using sparse canonical correlation analysis.

**Supplementary Table 3. Covariance between thalamic imaging metrics and performance in language processing**

| Modality | Thalamic imaging metrics | R |
|---|---|---|
| Triple-modality model | Controllability + Diffusion + Volume | 0.522 |
| Dual-modality model | Controllability + Diffusion | 0.503 |
| | Controllability + Volume | 0.397 |
| | Diffusion + Volume | 0.418 |
| Single-modality model | Controllability | 0.370 |
| | Diffusion | 0.394 |
| | Volume | 0.213 |

R values are generated using sparse canonical correlation analysis.

**Supplementary Table 4. Covariance between thalamic imaging metrics and performance in reasoning and encoding**

| Modality | Thalamic imaging metrics | R |
|---|---|---|
| Triple-modality model | Controllability + Diffusion + Volume | 0.487 |
| Dual-modality model | Controllability + Diffusion | 0.438 |
| | Controllability + Volume | 0.442 |
| | Diffusion + Volume | 0.349 |
| Single-modality model | Controllability | 0.371 |
| | Diffusion | 0.281 |
| | Volume | 0.286 |

R values are generated using sparse canonical correlation analysis.

**Supplementary Table 5. Covariance between thalamic imaging metrics and performance in visuospatial processing**

| Modality | Thalamic imaging metrics | R |
|---|---|---|
| Triple-modality model | Controllability + Diffusion + Volume | 0.503 |
| Dual-modality model | Controllability + Diffusion | 0.467 |
| | Controllability + Volume | 0.397 |
| | Diffusion + Volume | 0.354 |
| Single-modality model | Controllability | 0.354 |
| | Diffusion | 0.324 |
| | Volume | 0.207 |

R values are generated using sparse canonical correlation analysis.

**Supplementary Table 6. Covariance between thalamic imaging metrics and performance in impulse control**

| Modality | Thalamic imaging metrics | R |
|---|---|---|
| Triple-modality model | Controllability + Diffusion + Volume | 0.456 |
| Dual-modality model | Controllability + Diffusion | 0.429 |
| | Controllability + Volume | 0.418 |
| | Diffusion + Volume | 0.280 |
| Single-modality model | Controllability | 0.382 |
| | Diffusion | 0.230 |
| | Volume | 0.212 |

R values are generated using sparse canonical correlation analysis.

**Supplementary Table 7. Covariance between thalamic imaging metrics and performance in overall cognitive performance**

| Modality | Thalamic imaging metrics | R |
| --- | --- | --- |
| Triple-modality model | Controllability + Diffusion + Volume | 0.574 |
| Dual-modality model | Controllability + Diffusion | 0.565 |
| | Controllability + Volume | 0.515 |
| | Diffusion + Volume | 0.455 |
| Single-modality model | Controllability | 0.462 |
| | Diffusion | 0.446 |
| | Volume | 0.367 |

R values are generated using sparse canonical correlation analysis.
